# Suppression of Glucosylceramide Synthase Reverses Drug Resistance in Cancer Cells Harbor Homozygous p53 Mutants

**DOI:** 10.1101/2025.11.02.686136

**Authors:** Md Saqline Mostaq, Mohammad N Amin, Amanda Raphael, Celine Asbury, Anish Gupta, Xin Gu, Xianlin Han, Davorka Sekulic, Pawel Michalak, Lin Kang, Yong-Yu Liu

## Abstract

Glucosylceramide synthase (GCS) catalyzes ceramide glycosylation in response to cell stress that produces glucosylceramide and other glycosphingolipids. GCS overexpression is a cause of drug resistance and enriches cancer stem cells (CSCs) during cancer chemotherapy. Previous studies showed that GCS modulates expression of p53 mutants and oncogenic gain-of-function (GOF) in heterozygous knock-in cell models (*TP53* R273H^-/+^). However, it is unclear whether GCS can modulate the effects of homozygous p53 mutations, which are common in many cancer cases. We report herewith that inhibition of GCS, via UGCG-knockout and using new inhibitor (Genz-161), effectively re-sensitizes drug resistance and diminishes CSCs in colon cancer cells carrying the homozygous p53 R273H mutation. In aggressive WiDr cells carrying *TP53* R273H mutation, knockout of *UGCG* gene using CRISPR/Cas9 editing or inhibition of GCS with Genz-161 sensitized cancer cells to oxaliplatin, irinotecan and paclitaxel. With decreased ceramide glycosylation in lipidomic profiling, both UGCG-knockout and Genz-161 treatments substantially decreased wound healing, and diminished CSCs and tumor growth under chemotherapy. Interestingly, inhibition of RNA m^6^A methylation by neplanocin A reactivated p53 function and reversed drug resistance. Mechanistic investigation revealed that GCS inhibition downregulated METTL3 expression and repressed RNA-m^6^A modification on mutant p53 R273H effects. Altogether, our findings demonstrate that ceramide glycosylation promotes METTL3 expression and RNA m^6^A methylation in response to drug-induced stress, thereby promoting mutant p53 expression and associated GOF. Conversely, inhibition of GCS can diminish CSCs and drug resistance via reduction of m^6^A modification and reactivation of p53 function. GCS inhibition is an achievable approach for mutant cancer treatment.

## Introduction

Ceramide glycosylation catalyzed by glucosylceramide synthase (GCS) modulates cell processes in response to stress, including exposure to various therapeutic agents for cancer treatments [1, 2]. GCS is a rate-limiting enzyme for glycosphingolipid (GSL) synthesis, catalyzing ceramide (Cer) glycosylation that converts Cer into glucosylceramide (GlcCer) and provides a precursor for various GlcCer based glycosphingolipids (GSLs) [3, 4]. In addition to other activities, GSLs associated with membrane proteins in GSL-enriched microdomain (GEM) modulates the cellular effects of these proteins, such as Src family kinases [5, 6]. Previous reports indicate that globo-series GSLs, in particular globotriosylceramide (Gb3), play crucial roles in modulating the transcription of genes via cSrc and β-catenin signaling pathways [6–8]. Under treatments with anticancer drugs, increased Cer-glycosylation by GCS and cellular GSLs modulate GEMs so as to promote the expression of particular genes, including multidrug resistance gene (MDR1), fibroblast growth factor 2 (FGF-2) and even p53 mutants of cancer cells for avoiding drug-induced cell death [7–9].

Missense mutant proteins produced from the *TP53* gene lack the tumor suppressive activity of wild-type protein has and often exhibit oncogenic gain-of-function (GOF) [10–12]. Human *TP53* gene encodes p53 protein, an essential tumor suppressor that stabilizes the genome with respect to the propensity for neoplastic transformation in normal cells. Acting as a transcription factor with homo-tetramer, p53 promotes the expression of its responsive genes, including p21, Bax, Puma and others, whereby p53-dependent cell proliferation arrest or apoptosis is executed in response to genotoxic stress [10]. However, *TP53* mutations are frequently detected in cancers, particularly in more than 80% of metastatic cancers or recurred cancers, such as those of ovaries and colon [13, 14]. Missense mutations at codons 175, 248, and 273 constitute approximately 19% of all p53 genetic alterations, thus these codons are referred to as mutation hotspots [15–17]. In addition to other oncogenic effects on tumor progression, missense p53 mutants are causative of cancer drug resistance and CSCs [9, 18, 19]. Restoring the expression of wild-type p53 or reactivating p53 function has been shown to re-sensitize cancer cells carrying *TP53* mutations to anticancer treatments [11, 20–22].

Cell stress upon treatments with anticancer drugs can enhance Cer-glycosylation and up-regulate the expression of p53 mutants [6, 11, 12, 23]. Previous studies show inhibition of Cer-glycosylation catalyzed by GCS can reverse drug resistance and decrease CSC population by cancer cells with heterozygous *TP53* R273H (*TP53* R273H^+/-^) [6, 9, 11]. Homozygous p53 mutants are more commonly detected in cancer cases and are key players in cancer progression. To understand whether Cer glycosylation by GCS is associated with drug resistance and CSC enrichment of cancer cells carrying homozygous p53 mutations derived pathologically, rather than genetic editing, we studied WiDr cancer cells carrying homozygous *TP53* R273H mutation and examined the effects of new GCS inhibitor GENZ 667161 (Genz-161).

## Materials and Methods

### UGCG Knockout Cell Lines Development and Cell Culture

The human WiDr (missense mutation *TP53* R273H) colon cancer cell line was purchased from American Type Culture Collection (ATCC; Manassas, VA) [6, 24]. Its subline, WiDr/UGCG^-^ were generated by knock-out of human UGCG (NCBI gene ID:7357) via CRISP/cas9 gene editing by Gene Script (Piscataway, NJ). Briefly, target sites were located by a guidance RNA (gRNA) and transient transfection of RNP (GenScript CRISPR single-guide RNAs:Cas9). The endogenous UGCG gene was targeted and mutated, resulting in consequential reduction of the expression of GCS protein. A single isogenic knockout cell clone was identified by Sanger sequencing screening and defined as WiDr/UGCG^-^. Primers for UGCG gRNA T3 site (5’>3’): forward, AGAGTAGAGAAGCCAGCACGA; reverse, CTAGAACACAGCAGGTTCCCA. Cells of WiDr or WiDr/UGCG^-^ were cultured in ATCC-formulated EMEM containing 10% fetal bovine serum (FBS), 100 units/mL penicillin, 100 µg/mL streptomycin and 584 mg/L L-glutamine. SW48 (wild type TP53) colon cancer cells (from ATCC) were cultured in RPMI-1640 media containing 10% fetal bovine serum (FBS), 100 units/mL penicillin, 100 μg/ mL streptomycin and 584 mg/L l-glutamine. All cells were maintained in an incubator humidified with 95% air and 5% CO_2_ at 37°C.

### Cell viability assay

Cell viability was assessed using the CellTiter-Glo luminescent cell viability assay kit (Promega, Madison, WI), as described previously [11, 12]. Briefly, cells (4,000 cells/well) were grown in 96-well plates overnight and then switched to 5% FBS medium containing drugs for 72-h treatments. For combination treatment, cells were cultured in 10% FBS medium containing Genz-161 (4 μM) for 48 h in advance and placed into 96-well plates for overnight growth and co-cultured with drugs for an additional 72 h. Cell viability was assessed in a Synergy HT microplate reader (BioTek, Winnooski, VT, USA), following incubation with reagent of CellTiter-Glo Luminescent Cell Viability kit (Promega, Madison, WI). A new GCS inhibitor, Genz-161 (GENZ 667161, (*S*)-quinuclidin-3-yl(2-(2-(4-fluorophenyl)thiazol-4-yl)propan-2-yl carbamate) was kindly provided by Sanofi Genzyme (Framingham, MA) [25, 26]. Oxaliplatin, paclitaxel, irinotecan hydrochloride and 5-fluorouracil were purchased from Sigma-Aldrich (St. Louis, MO, USA). Neplanocin A (a potential m^6^A methylation inhibitor, NPC) was purchased from Cayman Chemical (Cayman Chemical, Ann Arbor, MI). DF-A7 (a specific degrader/inhibitor of m^6^A reader YTHDF2) was purchased from ProbeChem (Catalog#PC-22438, ProbeChem, Shanghai, China) [27].

### RT-qPCR analysis

This assessment was performed as described previously [12, 28]. Briefly, total RNA was extracted from cells or tissues using an SV Total RNA Isolation kit (Promega, Madison, WI). Equal amounts of RNA (100 ng/reaction) of samples were applied for assays using OneStep RT-PCR kit (Qiagen, Germantown, MD). Pairs of primers synthesized from IDT (Coralville, IA) for UGCG (forward, 5’-CTT GGT TCA CGG GCT GCC TTA C-3’; reverse, 5’-GAAACCAGTTACATTGGC AGAGAT-3’) were used in PCR amplification. The PCR amplification was performed in 35 cycles of denaturation at 94°C for 30 seconds, annealing at 55°C for 30 seconds, and extension at 72°C for 60 seconds. Glyceraldehyde-3-phosphate dehydrogenase (GAPDH), an endogenous control (forward, 5’-GTCTCCTCTGACTTCAACAGCG-3’; reverse, 5’-ACCACCCTGTTGCTGTAGCCAA-3’) was amplified.

### Tumor-sphere formation assay

It was conducted as previously described with minor modification [8, 29]. Briefly, cells of WiDr or WiDr/UGCG^-^ (passages 3-10; 500 cells) in 10% FBS EMEM medium were placed into 96-well ultra-low binding microplates and cultured for 6 days. Images of tumor spheres were captured using an EVOS FL cell imaging system with color CCD camera (100 x magnification; Thermo Fisher Scientific). Cell viability of tumor spheres was further assessed with CellTiter-Glo reagent, as described above (Cell viability assay).

### Wound healing assay

It was conducted as described previously [11, 30]. Briefly, cells (2×10^5^/well) were planted in 6-well microplates in 10% FBS EMEM medium with agents and the wounds were scratched with 100-μL pipette tips after 24 h growth (∼80% confluency). The wounded areas were observed and captured using EVOS FL cell imaging system (100 x magnification). For combination treatments, WiDr cells were pre-treated with Genz-161 (4 μM, 48 h) and then treated with IRI for an additional 48 h.

### Imaging flow cytometry analysis

Imaging flow cytometry was carried out as described previously [9, 11]. After treatments, suspended cells (10^6^ cells/ml) were incubated with human CD44v6 Alexa Fluor^®^ 488-conjugated antibody (2F10; mouse IgG1; from R&D Systems, Minneapolis, MN, USA) and human CD133 APC-conjugated antibody (170411; mouse IgG2b; from R&D Systems) in 1% BSA-containing PBS at 4°C for 45 min. After washing, cells were resuspended in 1% BSA PBS (5 x 10^5^ cells/150 μL) and analysed using an Amnis Imagestream Mark II Imagestream software, and the data were further analysed using the IDEAS v6.2 program.

### Western blot analysis

Western blotting was carried out as described previously [11, 12, 31]. Briefly, cells or tissue homogenates were lysed in NP40 cell lysis buffer containing protease inhibitors (Biosource, Camarillo, CA) to extract total cellular proteins following these treatments. Equal amounts of proteins (50 µg/lane) were resolved by using 4-12% gradient SDS-PAGE (Life Technologies). The nitrocellulose-membrane blots transferred were blocked in the StartingBlock Blocking buffer (proprietary protein formulation in 0.05% Tween-20, 20 mM phosphate-buffered saline, pH 7.5 (PBST), ThermFisher), and then incubated with each one of the primary antibodies (1:1000 or 1:2000 dilution), at 4 °C overnight. These blots were incubated with corresponding horseradish peroxidase–conjugated secondary antibodies (1:5,000 dilution) and detected using SuperSignal West Femto substrate (Thermo Fisher Scientific) and ChemiDoc MP imaging system (Bio-Rad, Hercules, CA, USA). Glyceraldehyde-3-phosphate dehydrogenase (GAPDH) was used as a loading control for cellular protein. Antibodies against human p53 phosphorylated at Ser15 were purchased from Cell Signaling Technology (Danvers, MA). Antibodies for p21, Bax, p53, and GAPDH were obtained from Santa Cruz Biotechnology (Dallas, TX). Relative protein levels were calculated from optical density values for each protein band using the Image Lab software v6.1 (Bio-Rad, Hercules, CA), normalized against those for GAPDH from three separate blots.

### Animal studies in tumor-bearing mice

All animal experiments were approved by the Institutional Animal Care and Use Committee, University of Louisiana at Monroe (ULM), and were carried out in strict accordance with good animal practice as defined by NIH guidelines. Athymic nude mice (Foxn1^nu^/Foxn1^+^, 7-9 weeks, female and males) were purchased from Envigo (Indianapolis, IN) and maintained in the vivarium at ULM. Animal studies were conducted as described previously [11, 12]. Briefly, cell suspension of WiDr/mock and WiDr/UGCG^-^ (5-7 passages, 2.5 × 10^6^ cells in 40 μL/each) was subcutaneously injected into the left flank or both left and right flanks of the mice. Mice with tumors (∼5 mm in diameter) were randomly allotted to different groups (6 mice/group, including 3 males and 3 females) for treatments. For treatments, Oxaliplatin (Oxa, 2.0, 4.0 mg/kg, once every 6 days) or irinotecan (IRI, 6 mg/kg) were administered intraperitoneally alone or with Genz-161 (4.0 mg/kg once every 3 days; in mice bearing WiDr tumors). Mice were monitored by measuring tumor sizes, body weights, and clinical observations, twice a week. Bone marrow were extracted and counted with hemocytometer as described previously [32]. Tumors and other tissues were dissected and stored at -80 °C for further analyses.

### Immunohistochemistry

Immunocytochemistry assessments were accomplished as described previously [11, 12, 31]. Microsections (5 μm) of tumors were prepared and stained with hematoxylin and eosin (H&E) by AML Laboratories (St. Augustine, FL) and further characterized by a pathologist. Antigens were retrieved in steaming sodium citrate buffers (10 nM, 0.05% Tween-20, pH 6.0). After blocking with 5% goat serum in PBS, slides were incubated with primary antibodies or conjugated antibodies (1:100) in blocking solution, at 4°C overnight. For detection of colon CSCs, the blocked slides were incubated with CD44v6 Alexa-Fluor 488-conjugated antibody (2F10; mouse IgG1; purchased from R&D Systems, Minneapolis, MN) and CD133/2 APC-conjugated antibody (293C3, mouse IgG2b; purchased from Miltenyi Biotec, San Diego, CA) (1:100). Corresponding Alexa Fluor 488-conjugated anti-mouse IgG and Alexa Fluor 594-conjugated anti-rabbit IgG (1:1000) were applied for further incubation to recognize the corresponding primary antibodies. After washing, cell nuclei were counterstained with DAPI (4, 6-diamidino-2-phenylindole) in mounting solution (Vector laboratories, Burlingame, CA, USA). Images (200× magnification) were captured using the EVOS FL cell imaging system with color CCD camera (Life Technologies, Grand Island, NY). Images (200× magnification) were captured using the EVOS FL cell imaging system with color CCD camera (Life Technologies, Grand Island, NY). Alexa Fluor 488-conjugated anti-mouse IgG and Alexa Fluor 594-conjugated anti-rabbit IgG were purchased from Thermo Fisher Scientific.

### Shotgun lipidomics analysis

Mass spectrometry-based shotgun lipidomics analysis was conducted, as described previously [33, 34]. Briefly, after treatments, tumor tissues were freshly dissected and stored at -80°C (5 mg each). Tissue samples were further homogenized in 0.1x PBS. Protein concentrations were measured using a bicinchoninic acid (BCA) assay prior to lipid extraction. A premixture of lipid internal standards including N18:0-d35 GalCer (20 nmol/mg proteins) for cerebroside quantitation was separately added to individual homogenate, based on the protein content of the sample. Lipid extraction was conducted by using a modified Bligh-Dyer, as previously described [35]. Each lipid extract was reconstituted with a volume of 500 μL/mg protein in CHCl_3_/MeOH (1:1, v/v). The lipid extracts were finally flushed with nitrogen, capped, and stored at -20°C for ESI/MS analyses.

ESI/MS quantitative analyses of cerebroside species were achieved using a triple-quadrupole mass spectrometer (TSQ Altis^TM^ mass spectrometer, Thermo Fisher ScientificSA) coupled with a Nanomate device (Advion, Ithaca, NY, USA) and controlled by the Xcalibur system [36]. Major cerebroside molecular species were directly quantitated by comparisons of ion peak intensities with that of an internal standard (*i.e*., N18:0-d35 GalCer) from the first-dimensional mass spectrum after correction for the different ^13^C-isotopologue intensities of cerebroside species relative to the internal standard in the positive-ion mode. By using these previously quantified major cerebroside species as a set of standards in addition to the original internal standard (*i.e*., N18:0-d35 GalCer), minor cerebroside species were quantitated/refined by MS/MS in a 2D mass spectrometric manner [36]. Data processing, including ion peak selection, baseline correction, data transfer, peak intensity comparison ^13^C deisotoping and quantitation, was facilitated by a custom-programmed Microsoft Excel macro as previously described [37, 38], ensuring accurate analysis of lipid molecular species. Quantitative data from biological samples were normalized to the protein content of the tissues, and all data are presented as means ± SD of samples.

### Data analysis

*Data analysis.* All experiments in cell models were performed in triplicate and repeated twice unless otherwise stated. Data are expressed as mean ± SD. One-way ANOVA with Tukey’s post-hoc test was applied to compare the continuous variables between groups. A two-way ANOVA with Bonferroni’s post-hoc test was applied among multiple comparisons (Figs. 1C, 4A), using the Prism v10 program (GraphPad, San Diego, CA). All p<0.01 comparisons were regarded as statistically significant.

**Figure 1.**
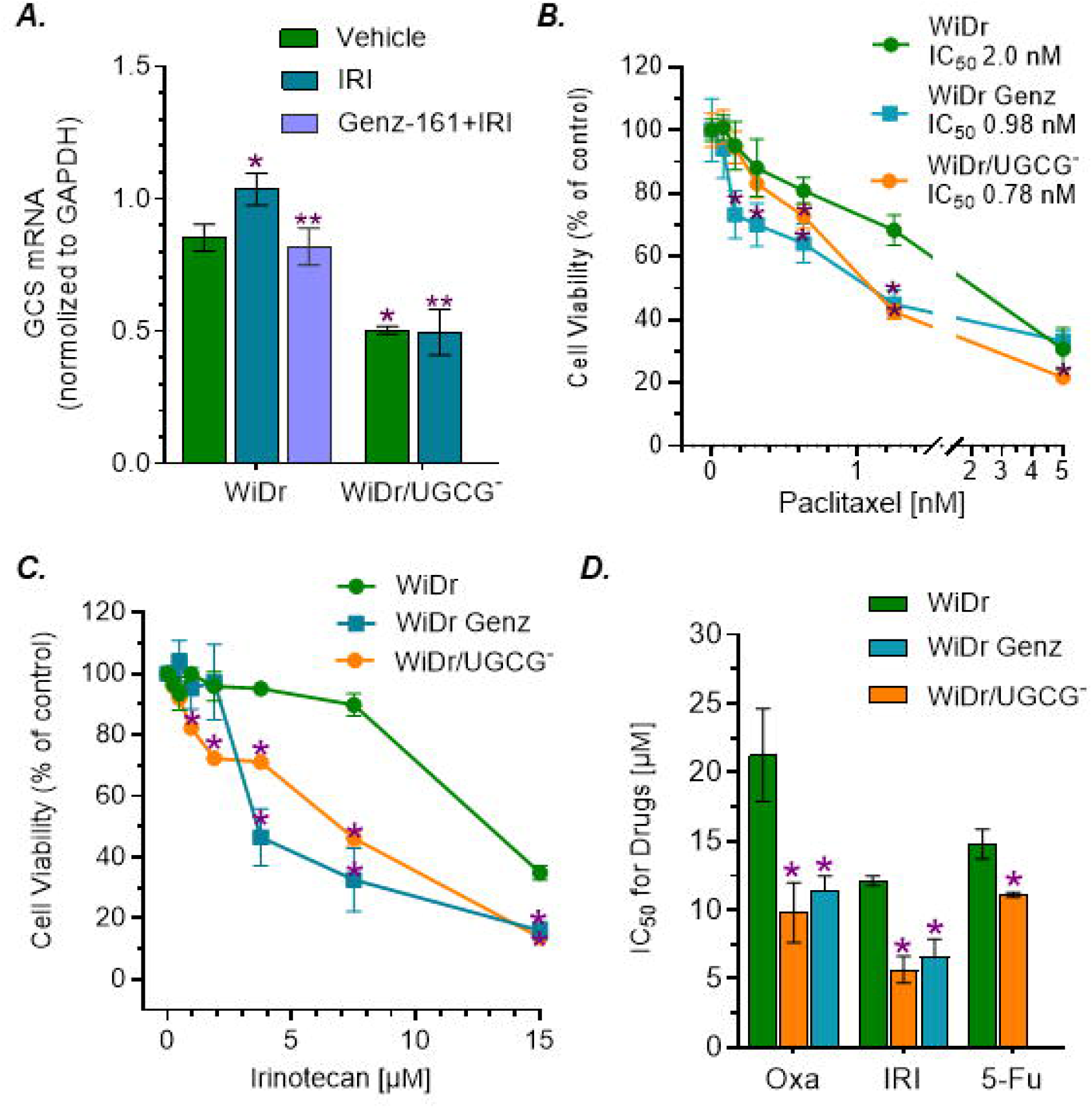
Effect of GCS on Drug Resistance in WiDr Cancer Cells Carrying *TP53* R273H. **A,** GCS mRNA levels in cancer cells. Cells were treated with 4 μM Genz-161 or vehicle for 48 h, and then exposed to irinotecan (IRI, 6 μM) for an additional 48 h. *, *p*<0.001 compared WiDr cells treated with vehicle control. **, p<0.001 compared with WiDr cells treated with IRI. **B,** Cell response to paclitaxel. **C,** Cell responses to irinotecan. WiDr cells were pretreated with 4 μM Genz-161 or vehicle for 48 h, and then co-treated with paclitaxel (Taxol) or irinotecan for an additional 72 h. *, *p*<0.001 compared with WiDr cells treated with vehicle. **D,** Effect of GCS inhibition on the IC_50_ values for anticancer drugs. *, *p*<0.001 compared with WiDr cells treated with vehicle.

## Results

### GCS Inhibition Reversed Drug Resistance and Reduced Tumorigenesis in p53 Mutant Cancer Cells

Parental WiDr cells, derived from human colorectal adenocarcinoma carrying homozygous *TP53* R273H mutation, displayed resistance to anticancer drugs, including doxorubicin (Dox), oxaliplatin (Oxa), 5-fluororacil (5-Fu) [6, 9, 39]. Following CRISPR/Cas9-edited UGCG-knockout, WiDr/UGCG^-^ cells reduced the expression of GCS mRNA (Fig. 1A). The levels of GCS mRNA were significantly decreased in WiDr/UGCG^-^ cells alone or treated with irinotecan (IRI), compared to WiDr cells or treated with IRI alone and combined with Genz-161 (Fig. 1A). We examined cell response to several drugs that often apply to chemotherapy of colorectal cancer, and found that WiDr/UGCG^-^ cells are sensitive to these drugs tested, including Oxa, 5-Fu, paclitaxel (Taxol) and irinotecan (IRI) (Fig. 1B-D). UGCG-knockout significantly increased the responses of WiDr/UGCG^-^ cells to treatments of Oxa, 5-Fu, Taxol and IRI (Fig 1B, 1C), and decreased the IC_50_ values by approximately 3-fold (0.78 nM *vs.* 2.0 nM, Fig. 1B) for Taxol, and 2-fold (5.7 μM *vs.* 12 μM, Fig. 1D) for IRI and Taxol, respectively. Genz-161 treatments that inhibit GCS enzyme activity, rather than decrease mRNA levels (Fig. 1A) [6], re-sensitized WiDr cells and reduced the IC_50_ values for Oxa, Taxol and IRI by approximately 2-fold (Fig 1B-D), respectively. Previous studies showed that GCS inhibition reversed Dox resistance in knock-in p53 mutant [11]. Present results indicate GCS is involved in modulating multidrug resistance of homozygous *TP53* R273H cancer cells, and further Genz-161, a new GCS inhibitor can effectively re-sensitize mutant cancer cells to anticancer drugs.

To further assess functional consequences of GCS inhibition, we conducted assays of tumorsphere formation and wound healing. Indeed, GCS inhibition decreased the tumorigenesis and cellular migration of WiDr cancer cells exposed to IRI. WiDr/UGCG^-^ cells formed significantly fewer and smaller tumorspheres, compared to WiDr cells (approximately 70% reduction; Fig. 2A, 2B). Genz-161 treatment displayed the same effects as UGCG-knockout, significantly decreasing tumorsphere formation of WiDr cells (Fig. 2A, 2B). IRI exposure enhanced the wound healing of WiDr cells, however, under the same condition, the wound healings were markedly decreased in either WiDr/UGCG^-^ cells or WiDr cells treated with Genz-161, compared with WiDr cells (Fig. 2C, 2D). At 24 h and 48 h post-scratch, the wound areas in WiDr/UGCG^-^ cells remained approximately 2-fold (44% *vs.* 78%) and 3-fold (13% *vs.* 49%) larger than these in WiDr cells (Fig. 2D). Genz-161 treatment displayed a similar effect on decreasing the wound healing of WiDr cells exposed to IRI.

**Figure 2.**
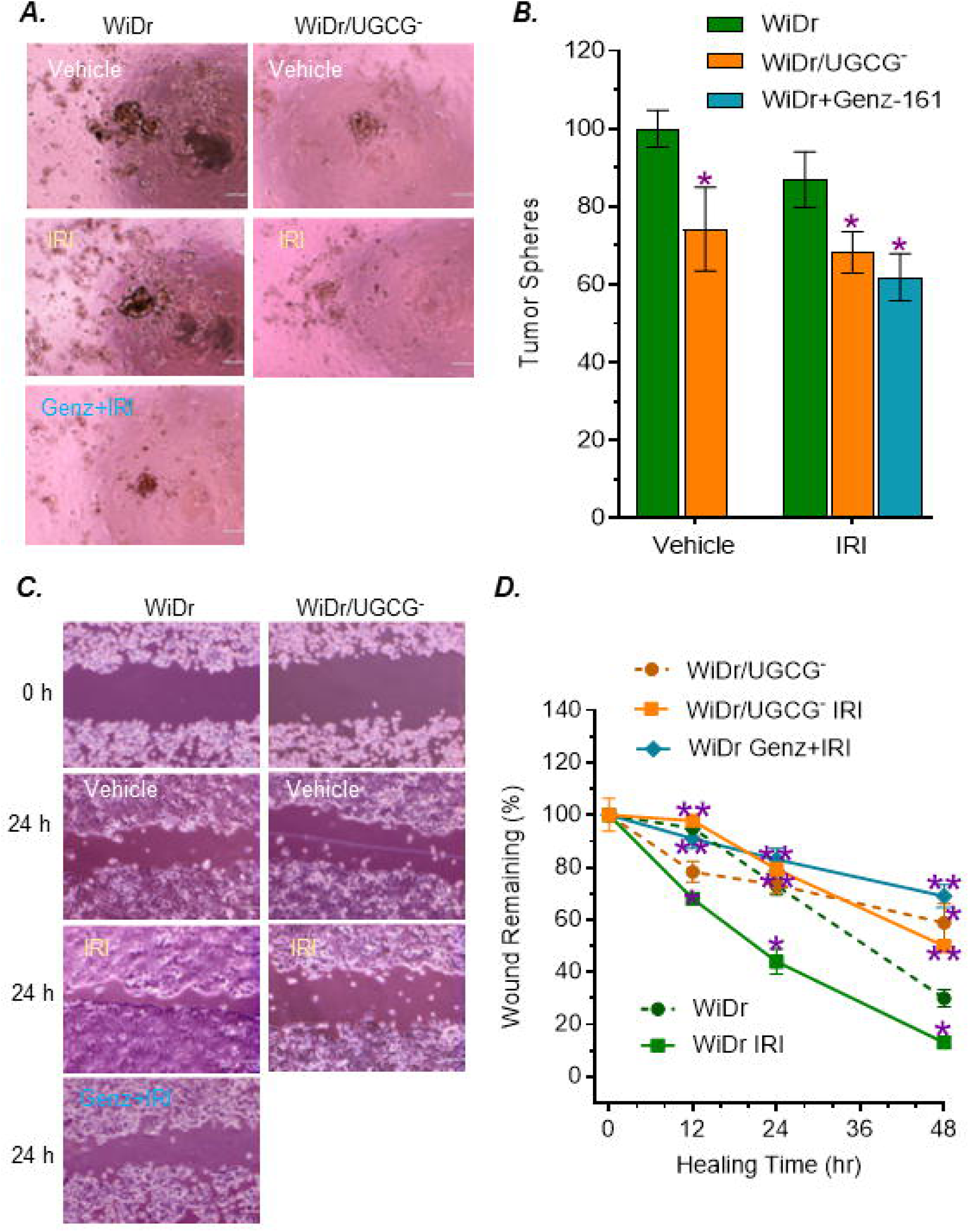
Inhibition of GCS Decreased Tumor Sphere and Wound Healing of WiDr Cancer Cells. **A,** Tumor sphere of WiDr cells. After pre-treatments of Genz-161 (4 μM) or vehicle, cells were planted in 5% FBS EMEM medium containing Genz-161 alone or combined irinotecan (IRI, 6 μM) for 6 days. Images were captured with 100x magnification. Scale bar represents 100 μm. **B,** Inhibition of GCS reduced tumor sphere formation. *, p<0.001 compared with WiDr cells treated with vehicle or IRI. **C**, Wound healing. Cells were pretreated with Genz-161 (4 μM) or vehicle for 48 h, and then exposed to IRI (6 μM) for an additional 48 h. WiDr/UGCG cells treated with IRI alone for 48 h. Images were magnified 100x, scale bar equals to 100 μm. **D**, GCS inhibition decreased the wound healing of WiDr cells. *, *p*<0.001 compared with WiDr cells treated with vehicle. **, *p*<0.001 compared with WiDr cells treated with IRI.

### GCS Inhibition Reduced the Enrichment of Cancer Stem Cells through Modulating the Expression of TP53 R273H Mutant Proteins

Previous studies indicate GCS can enrich CSCs during chemotherapy and contribute to drug resistance and metastasis [8, 11, 40]. Using imaging flow cytometry, we assessed cancer stem cells (CSCs) in WiDr cell lines exposed to IRI. In vehicle treatments, there is no significant difference between WiDr and WiDr/UGCG^-^ cells (Fig. 3A, 3B). Irinotecan treatments significantly increased CSC population in WiDr cells, compared to WiDr cell treated with vehicle (Fig. 3A, 3B). Further, we found that under irinotecan treatments, GCS inhibition significantly decreased CSC numbers of WiDr/UGCG^-^ cells or WiDr treated with Genz-161, by approximately 2-fold compared with WiDr cells (8.8% *vs.* 4.7% or 3.9%, *p*<0.01) (Fig. 3A-3C). These results indicate that anticancer drugs may enrich CSCs via ceramide glycosylation by GCS. Inhibition of GCS, by either genetic manipulation or enzyme inhibitor, can significantly decrease CSC enrichment during drug treatments.

**Figure 3.**
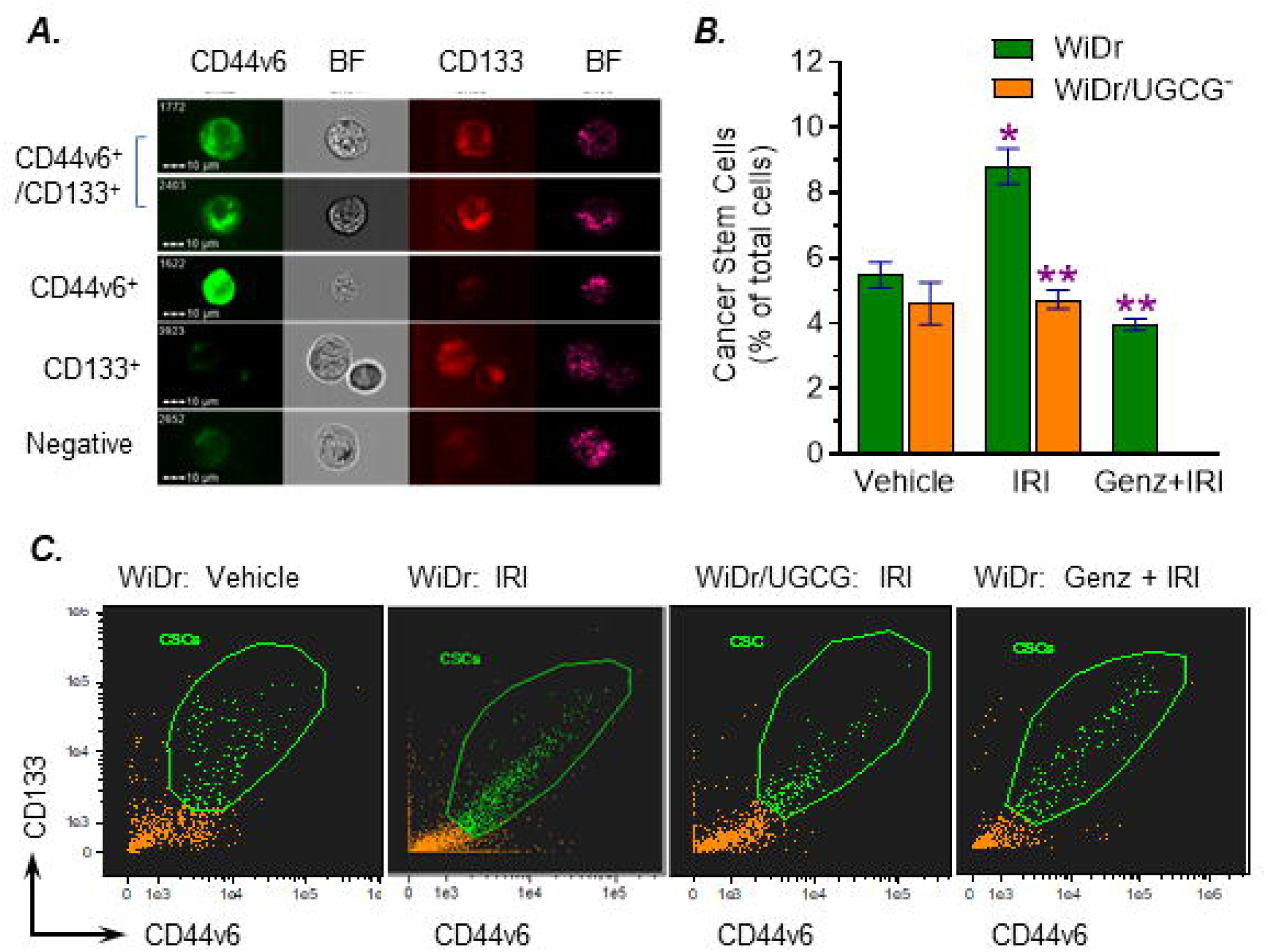
GCS Inhibition Decreased Cancer Stem Cell Population of WiDr Cells Exposed to Irinotecan. Cells were planted in 5% FBS EMEM medium containing 6 μM IRI alone or combined 4 μM Genz-161 for 6 days. **A,** Imaging flow cytometry analysis for colon CSCs (CD44v6^+^/CD133^+^). BF, bright field. **B,** GCS inhibition decreased CSC population. *, *p*<0.001 compared to WiDr cells treated with vehicle; **, *p*<0.001 compared to WiDr cells treated with IRI alone. **C,** Representative CSC plots of cancer cells with various treatments.

To understand how inhibition of GCS reverse drug resistance and tumorigenesis of WiDr cells carrying p53 mutation, we assessed the alterations of protein expression of cells under treatments. Irinotecan treatments significantly increased GCS protein levels of WiDr cells, compared with vehicle (Fig. 4A, 4B). With the lower mRNA levels (Fig. 1A), GCS protein levels were significantly decreased, by approximately 3-fold (0.24 *vs.* 0.69) and 6-fold (0.20 *vs.* 1.23, *p*<0.001) in WiDr/UGCG^-^ cells treated with vehicle or irinotecan, compared with these of WiDr cells (Fig. 4A, 4B). Genz-161 treatments did not have significant effect on GCS protein levels of WiDr cells (Fig. 4A, 4B). Interestingly, GCS inhibition substantially increased the levels of phosphorylated p53 (pp53, Ser15), by approximately 6-fold (0.13 *vs.* 0.82, *p*<0.001) in WiDr cells treated with Genz-161 and 8-fold (0.13 vs. 1.05, *p*<0.001) WiDr/UGCG^-^ cells treated with IRI (Fig. 4A, 4B). Consistently, the protein levels of p53-responsive genes, such as p21 and Bax that can induce cell proliferation arrest and apoptosis, are also significantly increased by approximately 2-fold (0.13 *vs.* 1.04, *p*<0.001) to 10-fold (0.05 *vs.* 0.63, *p*<0.001) in cells with GCS inhibition (Fig. 4A, 4B). We also found methyltransferase-like 3 (METTL3), which is reported responsible for RNA m^6^A methylation [9, 41], were significantly decreased by GCS inhibition (∼5-fold, *p*<0.001), either in WiDr cells treated with Genz-161 and IRI or WiDr/UGCG^-^ cells treated with IRI alone (Fig. 4A, 4B). These results clearly indicate that GCS inhibition can effectively reactivate pp53 expression and its dependent function in suppressing cancer stem cells and reverse drug resistance. This reactivating effect of GCS inhibition on functional p53 expression may be correlated to decrease METTL3 expression.

**Figure 4.**
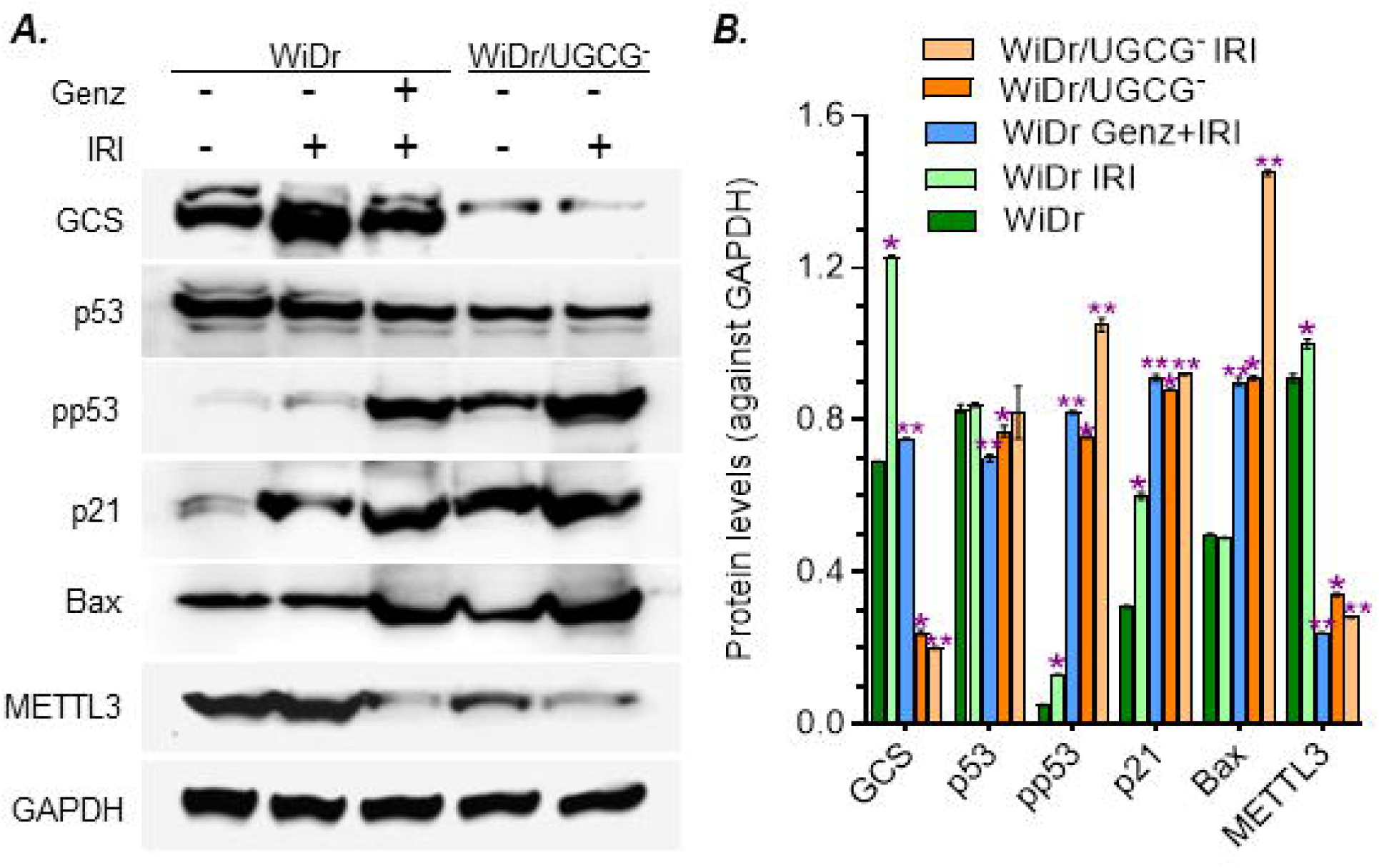
GCS Modulates Mutant p53 Protein Expression in WiDr Cancer Cells. **A,** Western blotting for protein expression. Cells were pretreated with 4 μM Genz-161 or vehicle for 48 h, and then exposed to 6 μM IRI for an additional 48 h. Equal amounts of soluble proteins (50 μg/lane) were resolved 4-12% SDS-PAGE and immunobloted with primary antibodies and HP-conjugated protein C. **B,** GCS modulated the expression of p53 R273H mutant. *, *p*<0.001 compared to WiDr cells treated with vehicle; **, *p*<0.001 compared to WiDr cells treated with IRI.

### GCS Inhibition Decreased Tumorigenesis and Sensitized Chemotherapeutic Effects in Mice-bearing p53 R273H Mutant Tumors

We examined the effects of GCS inhibition on tumorigenesis as well as drug resistance in mice bearing homozygous p53 R273H tumors. WiDr/UGCG^-^ tumors grew significantly slower and were more sensitive to Oxa treatments than WiDr tumors, consistent with decreased levels of GCS mRNA (Fig. 5A, 5D). In days 23-37 growth, tumor volumes of WiDr/UGCG^-^ are significantly decreased compared to tumor volumes of WiDr. After day 37 Oxa treatments, the volumes of WiDr/UGCG^-^ tumors (WiDr/UGCG^-^ Oxa) significantly decreased approximately 60%, compared with these of WiDr tumors (WiDr Oxa, Fig. 5A). Interestingly, Genz-161 treatments (4 mg/kg, per every 3 days) presented the same effect as UGCG-knockout did, these sensitized WiDr tumors to Oxa treatments and the tumor volumes of combination treatments (WiDr Genz+Oxa) decreased by approximately 60%, compared with WiDr treated with Oxa alone (WiDr Oxa) (Fig. 5A). Genz-161 also significantly sensitized WiDr tumors to IRI treatments (Fig. 5B). These treatments, including combination of Oxa and Genz-161, did not have significant effects on the body weights or other tissues (liver, kidney and bone marrow) (Suppl. Fig. S1).

**Figure 5.**
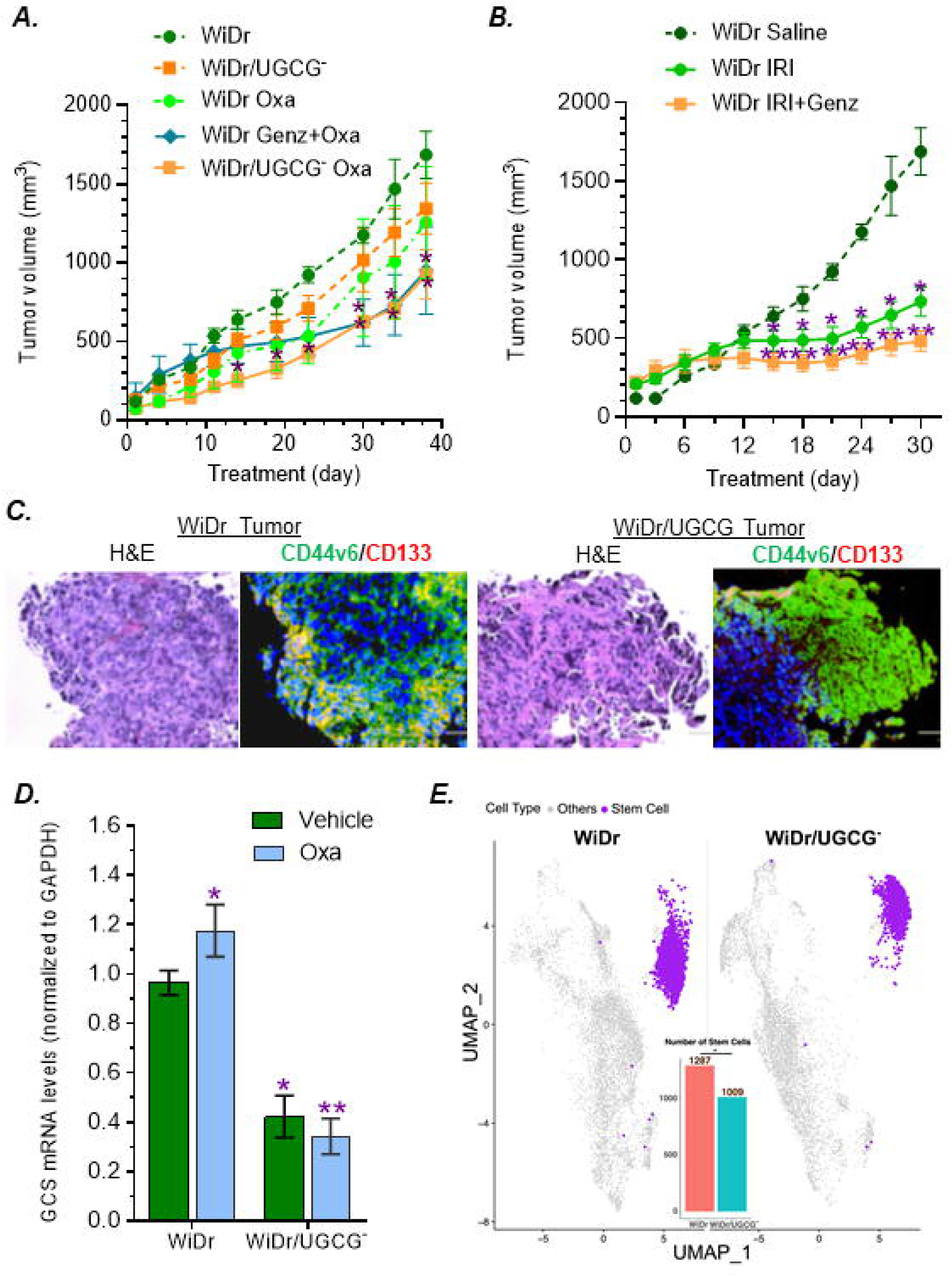
GCS Inhibition Decreased Tumor Growth. **A,** Tumor growth of mice treated with oxaliplatin. Mice-bearing tumors generated from WiDr or WiDr/UGCG^-^ cells were treated with vehicle, oxaliplatin (Oxa 2 mg/kg, *i.p*, once every 6 days) and combination (Oxa 2 mg/kg, i.p, once every 6 days and Genz-161 4 mg/kg, i.p, once every 3 days) for 37 days. *, *p*<0.01 compared with WiDr tumors treated with vehicle or Oxa. **B,** Tumor growth of mice treated with irenotecan. Mice-bearing tumors generated from WiDr cells were treated with vehicle, irenotecan (IRI 6 mg/kg, *i.p*, once every 6 days) and combination (IRI 6 mg/kg *i.p*, once every 6 days and Genz-161 4 mg/kg, i.p, once every 3 days) for 30 days. *, *p*<0.01 compared with tumors treated with vehicle; **, p<0.01 compared with tumors treated with IRI. **C,** H&E and immunofluorescence staining of tumors. Tumor sections were stained with H&E or fluorescent antibodies for CSC markers (CD44v6/CD133). Green, Alexa Fluor 448−CD44v6; red, APC-CD133; blue, DAPI nuclear counterstain. Images were magnified 200x, scale bar represents to 50 μm. **D,** GCS mRNA levels of tumors. *, *p*<0.01 compared with WiDr-tumors treated with vehicle or oxaliplatin. **E,** Tumor stem cell clusters in snRNA. Tumor-bearing mice were treated with Oxa (2 mg/kg, i.p, once every 6 days) for 37 days. *, *p*<0.001 compared with WiDr tumors treated with Oxa.

Immunofluorescence staining showed that colon CSCs (markers CD44v6^+^/CD133^+^) of WiDr/UGCG^-^ tumors significantly decreased by approximately 3-fold (9% *vs.* 3% of total tumor cells, *p*<0.001) compared with WiDr tumors treated with Oxa (Fig. 5C). Assessment of single-nuclear RNA-sequencing (snRNA-seq, detailed in our unpublished study) found that with decreased GCS mRNA levels, the stem cell cluster of WiDr/UGCG^-^ tumors is significantly decreased, compared with these of WiDr tumors treated with Oxa (Fig. 5E). snRNA-seq analysis further indicated differential expression of several key genes associated with CSC enrichment, drug resistance and tumor progression, including CD44, ZEB1, THSD4, RNF43 and DCBLD2 [11, 42–46], in p53 mutant tumors (Suppl Fig. S2).

### GCS Inhibition Modulated the Expression of p53 and p53-responsive Genes in Tumors Carrying p53 R273H Mutant

Previous works showed that inhibition of GCS increases cellular ceramide and decreases glucosylceramide, which can modulate mutant p53 expression [6, 22, 31]. We assessed and characterized the effects of inhibition of GCS on the expression of p53R273H in WiDr tumors. With increased levels of cellular ceramide stained with fluorescence-conjugative antibodies, nuclear pp53 protein levels are substantially upregulated in WiDr/UGCG^-^ tumors, compared with WiDr tumors treated with Oxa (Fig. 6A). Lipidomic analysis of tumor tissues further showed that total ceramide of WiDr/UGCG^-^ tumors are significantly increased with decreased cerebroside and increased sphingomyelin, compared with these of WiDr tumors treated with oxaliplatin (Fig. 6B, Supplementary S1). Among ceramide species, the levels of C16 ceramide (d18:1/16:0, N16:0, Fig. 6B) and C16 sphingomyelin (d18:1/16:0, N16:0, Suppl Fig. S3) are remarkably enhanced in WiDr/UGCG^-^ tumors, compared to these of WiDr tumors.

**Figure 6.**
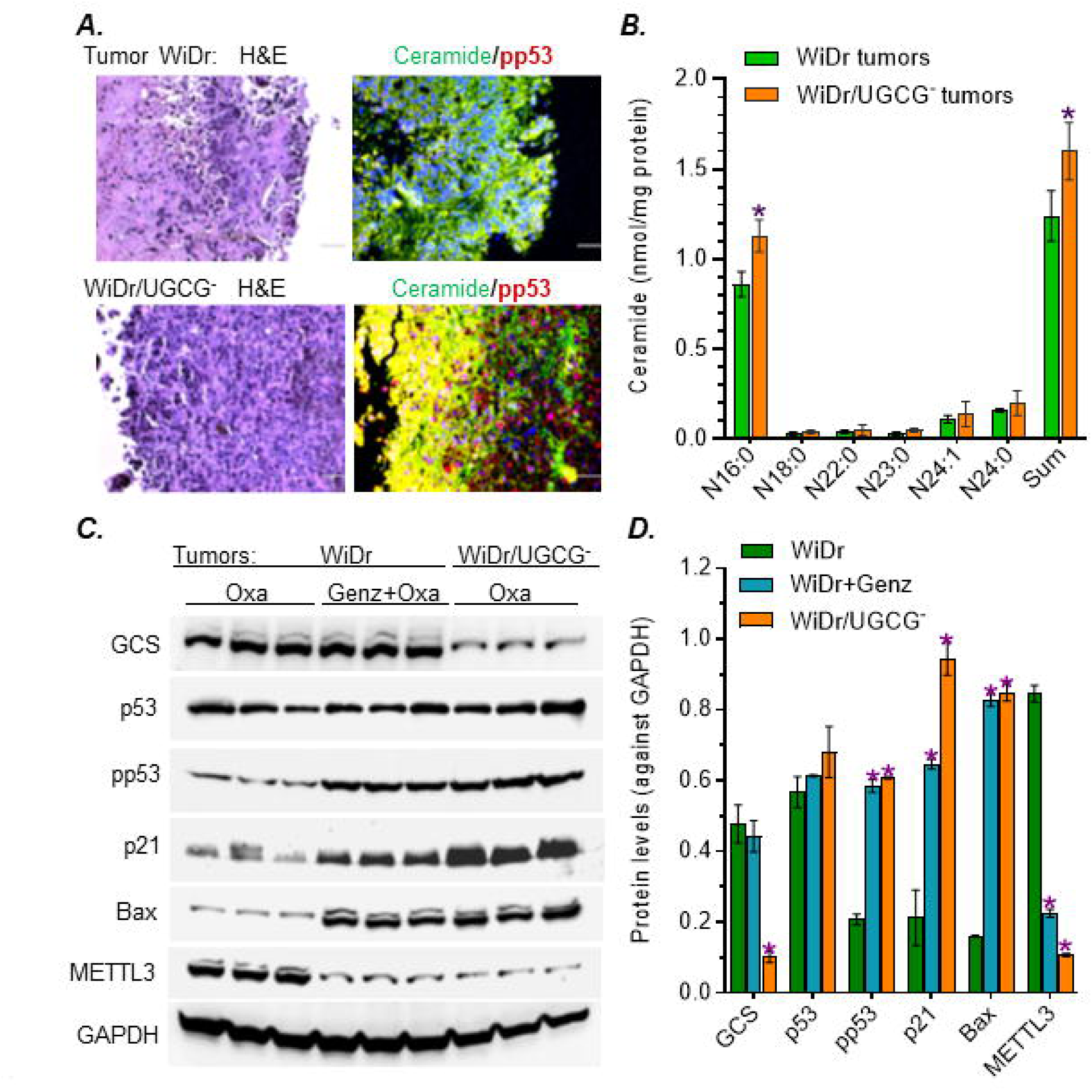
GCS Inhibition Sensitized Tumor Response to Chemotherapy. Tumor-bearing mice were treated with Oxa (2 mg/kg) alone or combined Genz-161 (Oxa 2 mg/kg and Genz-161 4 mg/kg). **A,** Immunofluorescence staining of tumors. Tumor sections were stained with H&E or fluorescent antibodies. Green, Alexa Fluor 488−ceramide; red, Alexa Fluor 555−pp53; blue, DAPI nuclear counterstain. Images were magnified 200x, scale bar represents to 50 μm. **B,** Lipidomics of tumors. *, *p*<0.001 compared with WiDr-tumors treated with Oxa. **C**, Western blotting of tumors. Tumors were pretreated with Oxa alone or combined with Genz-161. Equal amount of soluble proteins (50 μg/lane) were applied for Western blotting. **D,** GCS inhibition modulates mutant p53 protein expression of tumors. *, *p*<0.001 compared with WiDr-tumors treated with Oxa.

Consistent with the cell models (Fig. 4), we see GCS protein levels decreased by UGCG-knockout in WiDr/UGCG^-^ tumors treated with Oxa, compared with these of WiDr tumors (Fig. 6C, 6D). The protein levels of GCS in WiDr/UGCG^-^ tumors were decreased, by approximately 5-fold (0.1 *vs.* 0.48, *p*<0.001), compared to these of WiDr tumors with the same treatments (Fig. 6C, 6D). Combination treatments (Genz-161+Oxa) did not have any significant influence in GCS expression of WiDr tumors. Inhibition of GCS, by using either UGCG-knockout or Genz-161 treatments, reactivate functional p53 and p53-responsive genes to suppress tumor. The pp53 levels increased by approximately 3-fold (0.2 *vs.* 0.61, *p*<0.001) in WiDr/UGCG^-^ tumor and WiDr tumor treated with Genz-161, compared with WiDr treated with Oxa alone (Fig. 6C, 6D). Consequently, the protein levels of p21 and Bax are significantly increased by approximately 4-fold (0.21 *vs.* 0.94, *p*<0.001) and 5-fold (0.16 *vs*. 0.85, *p*<0.001) in WiDr/UGCG^-^ tumors, compared with WiDr tumors (Fig. 6C, 6D). Genz-161 treatments displayed similar effects in WiDr tumors treated with combination as UGCG-knockout did. METTL3 protein levels are significantly decreased by GCS inhibition, either in WiDr tumors treated with Genz-161 (4-fold, 0.23 *vs.* 0.81, *p*<0.001) or WiDr/UGCG^-^ tumors (7-fold, 0.11 *vs.* 0.81, *p*<0.001), compared with WiDr tumors treated with Oxa (Fig. 6C, 6D). These results further indicate that inhibition of GCS can reactivate the expression of pp53 and p53-responsive genes to suppress tumorigenesis and reverse drug resistance through modulation of RNA m^6^A methylation.

### Ceramide Glycosylation Upregulated the Expression of METTL3 and RNA m^6^A Methylation for Expressing p53 R273H Protein and Drug Resistance

We characterized the influence of m^6^A methylation in drug resistance of WiDr cells carrying the R273H *TP53* mutation, compared to SW48 cancer cells that carry wild-type *TP53* [11, 47]. Previous studies showed that neplanocin A (NPC) can inhibit RNA m^6^A methylation [9, 48]. We examined the m^6^A-RNA levels of cancer cells with treatments using ELISA. Cell stress of IRI treatments greatly increased the m^6^A-RNA levels of WiDr cells, by 2-fold (0.018% *vs.* 0.035%, *p*<0.001), compared with SW48 cells (Fig. 7A). NPC treatments significantly decreased m^6^A-RNA levels, by 1.6-fold (0.011% *vs.* 0.018%, *p*<0.001) in SW48 cell and by 2-fold (0.016% *vs*. 0.038%, *p*<0.001) in WiDr cells, respectively compared with cells exposed to IRI (Fig. 7A). NPC treatments significantly sensitized WiDr cells to IRI and significantly decreased the IC_50_ values, by approximately 4-fold (3.1 μM *vs*. 12 μM, *p*<0.001), compared with WiDr cells with vehicle (Fig. 7B). However, NPC treatments displayed less effect on the response to IRI of SW48 that carry wild-type *TP53* (Fig. 7B). In Western blot analysis, we found that METTL3 protein levels were higher in WiDr cells treated with either IRI alone or combination of IRI and NPC, by approximately 2-fold (0.71 *vs.* 1.4, *p*<0.001), compared with SW48 cells in the same conditions. NPC treatments decreased the protein levels of METTL3 in both WiDr and SW48 cell lines, by approximately 30%, compared to these treated with IRI alone. NPC treatments remarkably increased pp53 protein levels in WiDr cells treated with combination, by approximately 10-fold (0.13 *vs.* 1.3, *p*<0.001), compared with WiDr cells with IRI alone (Fig 7C, 7D). In consistency, the protein levels of p21 and Bax are also significantly increased in WiDr cells treated with NPC, by approximately 5-fold (0.21 *vs.* 1.13, *p*<0.001) and 4-fold (0.37 *vs*. 1.34, *p*<0.001) in WiDr cells treated with combination, compared with WiDr cells treated with IRI alone respectively (Fig. 6C, 6D). In the same treatment conditions, there is no significant difference the pp53 levels of SW48 cells treated with NPC compared to vehicle. These results clearly indicate that RNA m^6^A methylation catalyzed by METTL3 plays a critical role in modulating the expression of mutant p53 by cancer cells in response to drug-induced cell stress (Fig. 8).

**Figure 7.**
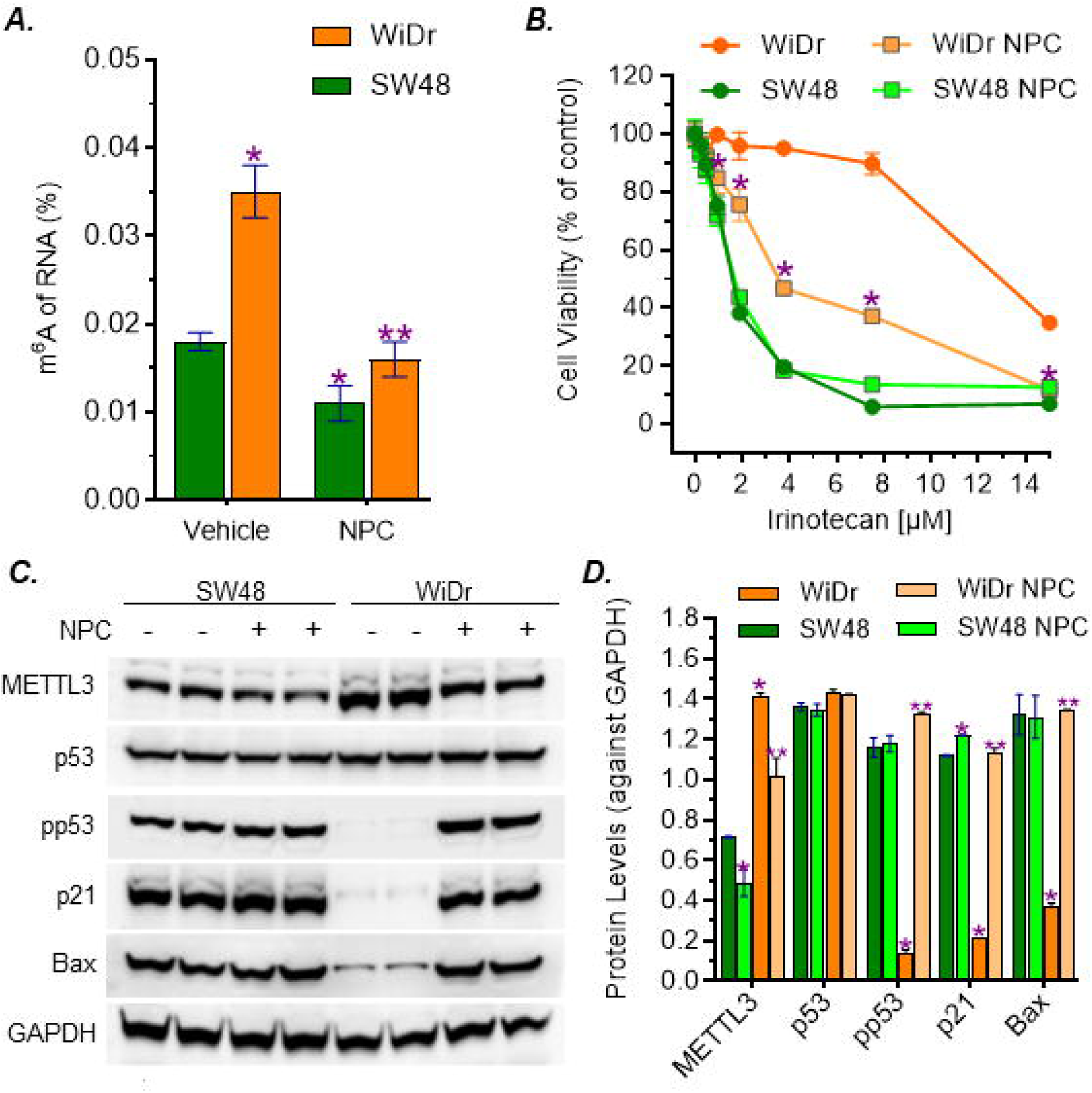
GCS Inhibiton Modulates the Expression of Mutant p53 R273H via RNA m^6^A Methylation. **A**, ELISA of RNA-m^6^A. Cells were pretreated with 200 nM NPC or vehicle for 48 h and co-treated with IRI for additional 48 h. Equal amount of extracted toral RNA (200 ng/reaction) was applied to ELISA. *, *p*<0.001 compared with SW48 cells treated with vehicle. **, p<0.001 compared with WiDr cell treated with vehicle. **B**, Cell response to IRI. Cells were pretreated with 200 nM NPC or vehicle for 48 h, and co-treated with increasing concentrations of IRI for additional 72 h. *, *p*<0.001 compared with WiDr cells treated with vehicle. IC_50_ values: WiDr = 14.79 ± 1.08 μM, WiDr NPC = 3.1 ± 0.05 μM (*p*<0.001 compared with WiDr treated with vehicle); SW48 = 1.45 ± 0.04 μM, SW48 NPC = 1.31 ± 0.03 μM. **C**, Western blots of METTL3 and p53 proteins. Cells were pretreated with 200 nM NPC or vehicle for 48 h and co-treated with 3 μM IRI for additional 48 h. Equal amount of soluble proteins (50 μg/lane) were applied for Western blotting. **D**. NPC treatments reactivate the expression of pp53 protein. *, *p*<0.001 compared with SW48 cells with IRI. **, p<0.001 compared with WiDr cells treated with IRI.

**Figure 8.**
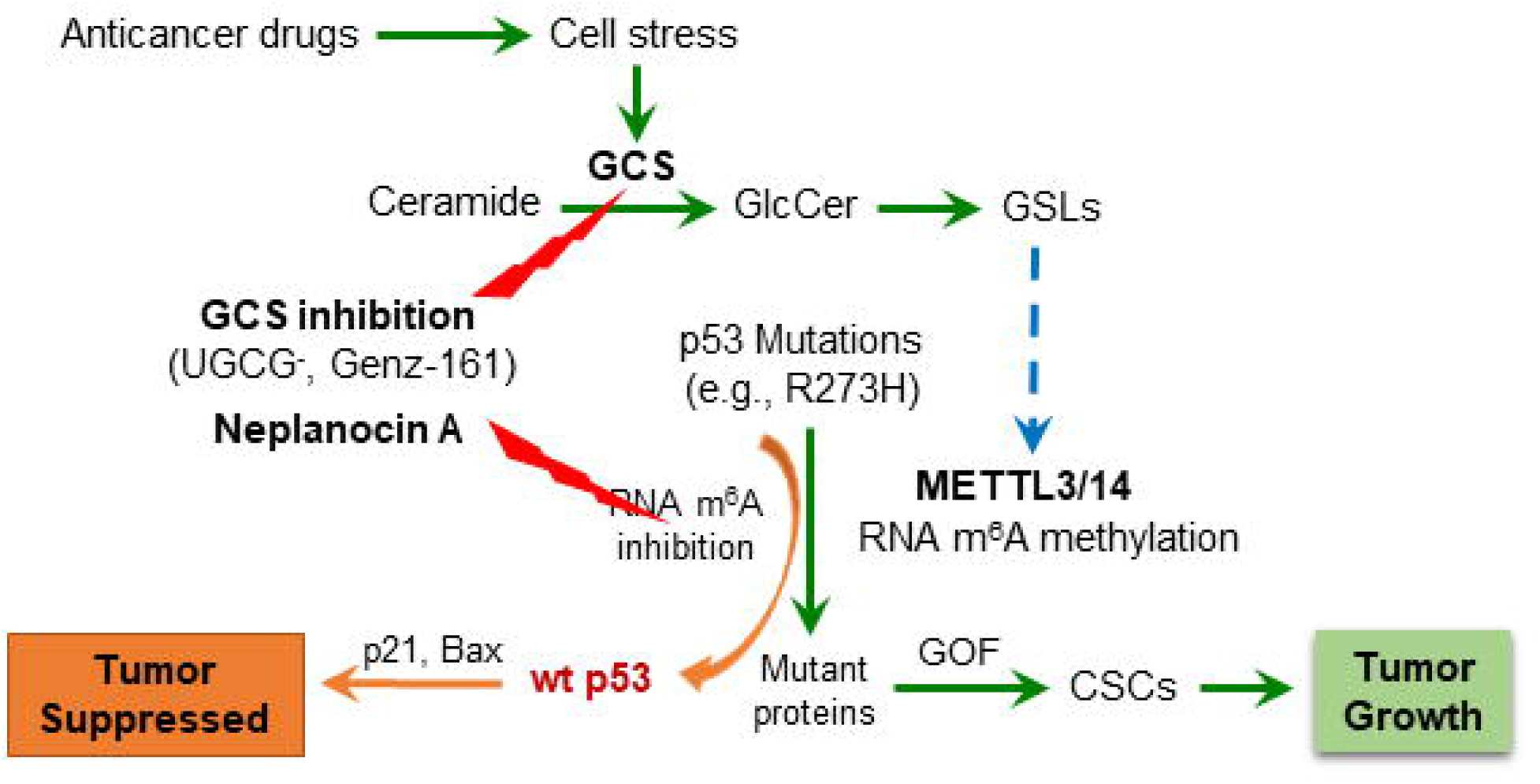
Ceramide Glycosylation by GCS Modulates RNA m^6^A Methylation in Response to Cell Stress Expressing Mutant p53 R273H. In response to anticancer drugs, cell stress upregulates the expression of glucosylceramide synthase (GCS) and ceramide glycosylation by cancer cells that enhances the production of glucosylceramide (GlcCer) and then glycosphingolipids (GSLs). GSLs induce expression of methyltransferase like-3 (METTL3) and RNA *N*^6^-methyladenosine (m^6^A) methylation of cancer cells carrying p53 mutations (*TP53* R273H). RNA m^6^A modification upregulates missense mutatant proteins and the gain-of-function (GOF) enriches cancer stem cells (CSCs) for tumor growth. Conversely, inhibition GCS (using either UGCG-knockout or inhibitor Genz-161) or supression of METTL3 and RNA m^6^A modification can effectively restore the expression of wild-type p53 protein (wt p53) and its function in tumor suppression.

## Discussion

Our present study indicates that GCS inhibition can reverse the drug resistance and inhibit the tumor progression of aggressive cancer cells carrying homozygous p53 R273H mutation. As common cell responsiveness to cell stress induced by anticancer drugs, Cer-glycosylation by GCS increases GSLs, including globotriosylceramide Gb3 [8, 49–51]. Overexpression of GCS and other enzymes in GSL synthesis are correlated with drug resistance and CSC of various cancers, including colorectal cancer, triple-negative breast cancer, hepatocarcinomatous and melanoma [52–54]. Targeting GCS as well as GSLs (GD2, Gb3) is emerging as an effective approach for improving the outcome of chemotherapy and immunotherapy [1, 2, 55–57]. Our previous works reveals that Cer-glycosylation by GCS can modulate the protein expression of p53 mutations and the oncogenic GOF of heterozygous p53 mutant models (p53 R273H^-/+^ generated by CRISP/cas9 editing) [6, 9, 11]. Our present studies demonstrate that inhibition of GCS, by either UGCG-knockout or GCS inhibitor, can reverse cancer drug resistance and eliminate CSC enrichment of homozygous p53 R273H cancer cells. WiDr cells, derived from colon adenocarcinoma carrying p53 R273H mutation, are notorious for drug resistance and aggressive growth [6, 58, 59]. In the present study, we see knockout UGCG gene that encodes GCS greatly re-sensitized WiDr/UGCG^-^ cells to several anticancer drugs, including IRI, Oxa, 5-Fu and Taxol, compared to parental WiDr cells (Fig. 1). In *vivo*, WiDr/UGCG^-^ tumors are sensitive to Oxa treatments, which is one of the first-line drugs used for colorectal cancer (Fig. 5). This specific approach for targeting UGCG gene immensely decreases the tumorigenesis and cell migration of WiDr/UGCG^-^ cells in tumorsphere formation, wound healing and tumor growth of mice (Fig. 2, Fig. 5). Indeed, UGCG-knockout eliminates the enrichment of colon CSCs in WiDr/UGCG^-^ cells and tumors (Fig. 3, Fig. 5C, 5D). These effects of UGCG-knockout on WiDr/UGCG^-^ cells are further supported by the evidence in treatments of Genz-161, a novel GCS inhibitor [6, 60]. Lower doses of Genz-161, either in cell models (4 μM) or in tumor-bearing mice (4 mg/kg, *i.p.*, per every 3 days), sufficiently reverse drug resistance and decrease tumor growth without significant adverse effects (Fig. 1, Fig. 5A, Suppl. Fig. S1). These results not only indicate that the key role played by GCS in tumor progression, but also indicate Genz-161 has a potential to be a potent therapeutic agent for cancers harbor homozygous p53 mutations, at least p53 R273H.

Suppressing ceramide glycosylation by GCS provides an effective option for targeting p53 mutant cancers. Targeting p53 mutations, particular missense mutations that are frequently detected in many types of cancer and promote tumor progression, is crucial for improving the outcome of cancer treatments [16, 20, 61–63]. Among these available options, restoration expression of wild-type p53 protein, which can eliminate the dominant effects of missense mutant p53 proteins included in the hetero-tetramers by cancer cells, might be more effective but face a great challenge [11]. Intriguingly, our previous studies elucidated that inhibition of Cer-glycosylation restores functional p53 tumor suppression, in heterozygous models (p53 R273H^-/+^) generated by knock-in editing [6, 9, 11]. Furthermore, the present study demonstrates that inhibition of GCS reactivates p53-dependent tumor suppression in WiDr colon cancer cells carrying homozygous mutation (p53 R273H) (Fig. 4, 6). Congruous with the effects of reversing drug resistance and tumorigenesis, UGCG-knockout or GCS inhibitor Genz-161 strikingly upregulated the protein levels of pp53 with p53-responsive genes (p21 and Bax) by homozygous p53-mutant WiDr cells, either cell culture with IRI or tumor-bearing mice treated with Oxa (Fig. 4, Fig. 6C, 6D).

The RNA m^6^A modification is involved in modulating the expression of p53 mutant proteins. The RNA m^6^A methylation that is mainly catalyzed by METTL3 in human cells can manipulate the fate of mRNA for producing protein in pre-mRNA splicing, mRNA stabilization and nuclear export for translation [9, 64, 65]. RNA m^6^A methylation and further modification with m^6^A readers, a groups of proteins binding to m^6^A-RNA, are correlated the expression of p53 mutant and neoplastic transformation [9, 66, 67]. Enhanced GSLs, particular Gb3 in GSL-enriched microdomain, can increase the expression of METTL3 via cSrc and β-catenin signal transduction pathway [6, 9]. Our previous studies showed that RNA m^6^A methylation by METTL3 at the mutant codon (R273H) regulated the expression mutant protein and GOF by SW48/TP53 Dox cells (*TP53* R273^-/+^) [9]. In current study, we found drug-induced cell stress significantly enhanced METTL3 protein levels in WiDr cancer cells or tumors (Fig. 4, Fig 6C-D). Conversely, inhibition of GCS, using either UGCG-knockout or Genz-161 markedly eliminated METTL3 protein expression in cell models and xenograft tumors (Fig. 4, Fig. 6C-D). Inhibition of RNA m^6^A methylation using NPC (Fig. 7A) inspiringly restored the expression of pp53 and of p53-responsive genes (p21, Bax) in WiDr cells carrying *TP53* R273H, rather than SW48 cells (Fig. 7C-D). NPC treatments significantly re-sensitized WiDr cell response to IRI (Fig. 7B), further indicating inhibition of RNA m^6^A methylation can reactivate tumor suppression in p53 mutant status. Currently, understanding of the mechanisms underlying RNA m^6^A modification and mutant protein expression is limited, despite their importance for improving cancer treatments.

Altogether, our present study demonstrated that Cer-glycosylation via RNA m^6^A methylation upregulates expression of p53 R273H and its GOF by cancer cells in response to chemotherapy. Conversely, suppression of Cer-glycosylation with a GCS inhibitor, or directly repressing m^6^A modification, can restore wild-type p53 function and re-sensitize drug-resistant cancerous cells and tumors to chemotherapeutic agents. Supported with our findings, inhibition Cer-glycosylation by GCS emerges as achievably therapeutic approach for targeting particular gene mutations to improve cancer treatments.

## Acknowledgments

We thank Katarzyna Michalak in Biomedical Affairs and Research, Edward Via College of Osteopathic Medicine for technical assistance.

## Authors’ contributions

Y.L., S. M. and L.K., designed research; S.M., M. A., A.R., C.A., A.G., X.G, X.H, Y. L. and L.K performed experiments and analyzed data; Y.L, L.K and P.M. supervised this work; S.M., Y.L. and L.K. wrote the main manuscript text and prepared figures; D.S contributed new reagents. All authors reviewed the manuscript.

## Data availability

The lipidomic data are publicly available within the article and its supplementary data files.

## Declarations

### Ethics approval and consent to participate

All animal experiments were approved by the Institutional Animal Care and Use Committee, University of Louisiana at Monroe (AWA #A3641-01) and were carried out in strict accordance with good animal practice as defined by NIH guidelines.

### Consent for publish

All the authors provided consent for the publication of the manuscript in the journal *Cancer Drug Resistance*

